# Fourfold lifespan extension in *C. elegans* daf-2 mutant males

**DOI:** 10.64898/2025.12.02.691904

**Authors:** Mike Russell, Michelle Lin, Alexander Tate Lasher, Steven N Austad, Liou Y Sun

## Abstract

Aging is a universal biological process driven by conserved genetic networks that balance somatic maintenance with growth and reproduction. The insulin/insulin-like growth factor I signaling pathway is a central architect of this balance, and inhibiting its activity has long been established as a primary mechanism to extend lifespan across diverse species. However, our understanding of this pathway’s limits has been constrained by a historical focus on hermaphroditic models, which inherently link longevity extensions to a rigid trade-off involving immense reproductive costs and restricted somatic growth. Here we show that the latent potential of this canonical aging pathway is profoundly amplified by male-specific biology. We demonstrate that a classic insulin receptor mutation extends male survival to an unprecedented extreme, vastly surpassing the established benchmarks of the field. We reveal that this extraordinary lifespan expansion remains strictly dependent on the canonical FOXO transcription factor, yet it fuels a male-specific metabolic reprogramming that uncouples aging from stunted growth—massively accumulating neutral lipids to support sustained somatic preservation. These findings establish biological sex as a primary determinant of longevity potential and provide a new framework for identifying hidden, sex-specific mechanisms capable of promoting extreme healthy longevity.

## Introduction

Aging is a universal biological process and is among the greatest risk factors for chronic disease (MacNee et al., 2014). While decades of research have uncovered conserved genetic pathways that regulate lifespan, the mechanisms that determine why individuals age differently remain unresolved. Among these regulators, the insulin/insulin-like growth factor I (IGF-I) signaling (IIS) pathway was the first discovered and remains one of the most powerful central determinants of longevity (Kenyon, 2011). Initial studies from Cynthia Kenyon and colleagues (1993) demonstrated that a single point mutation in the gene coding for DAF-2, a *Caenorhabditis elegans* homolog of the mammalian insulin or IGF-I receptor (Kimura et al., 1997), effectively doubled lifespan. Follow up studies in *Drosophila* (Clancy et al., 2001; Tatar et al., 2001) and mice (Bokov et al., 2011; Holzenberger et al., 2003) have also reported lifespan extension in mutants with interruptions along the IIS pathway. Further, mutations in the human *IGF1R* gene conferring reduced IGF-IR function have been identified in families with exceptional longevity (Suh et al., 2008). Together these position the IIS pathway as a potent, evolutionarily conserved mediator of longevity.

Almost as common as the lifespan extensions in IIS mutants are sex-specific differences in their lifespans. In the *Drosophila* and mouse studies above, significant longevity benefits were only achieved in one sex. Despite the commonality of these observations, very little work has been carried out on the influence of sex on IIS mutations in *C. elegans*. Worm studies largely employ hermaphrodites, leaving the impact of sex unstudied. This omission overlooks evidence suggesting that sex is a fundamental regulator of physiology. Profound differences in metabolism, reproductive strategy, and stress response between the sexes could fundamentally alter the output of core aging pathways (Austad & Fischer, 2016). Here, we investigate the influence of biological sex on the well documented *daf-2* mutant *C. elegans* model of longevity. We show that while *daf-2* hermaphrodites exhibit the canonical twofold lifespan extension, *daf-2* males experience a fourfold increase, coupled with a similar extension of healthspan. This work reveals that sex-specific biology can dramatically amplify the output of a conserved aging pathway, establishing sex as a primary determinant of longevity potential.

## Results

### Male daf-2 Mutants Live Twice as Long as Hermaphrodite daf-2 Mutants

To determine if the *daf-2(e1370)* mutation’s effect on lifespan was dependent on sex, we carried out three independently replicated lifespan analyses. Hermaphrodite *daf-2* mutants displayed notable extensions in lifespan over hermaphrodite WTs, consistent with previous reports (Kenyon et al., 1993). Surprisingly, male *daf-2* mutants markedly outlived hermaphrodite WTs, male WTs, and even hermaphrodite *daf-2* mutants (Fig. 1A-C). Hermaphrodite *daf-2* median lifespan (14.5 days) was comparable to that of the hermaphrodite WTs (14 days), however mean lifespan was 153.1% of the WT mean lifespan (14.7 days vs 22.5 days). Male *daf-2* median lifespan was 408.3% of the WT male median (18 days vs 73.5 days), and mean lifespan was 360.3% of the WT males (19.4 days vs 69.9 days). The male WT median lifespan was 128.6% of the hermaphrodite WT median (18 days vs 14 days), and the male WT mean lifespan was 132% of the hermaphrodite WT mean (19.4 days vs 14.7 days). Male *daf-2* median lifespan was a marked 506.9% of the hermaphrodite *daf-2* median (73.5 days vs 14.5 days) and male *daf-2* mean lifespan was 310.7% of the hermaphrodite *daf-2* mean (69.9 days vs 22.5 days). Hermaphrodite *daf-2* overall survival was significantly improved over hermaphrodite WTs (logrank P<0.0001), and hazard ratio was significantly reduced (HR=0.4753; P<0.0001). Survival at the 75^th^ and 90^th^ percentiles of life were significantly improved in hermaphrodite *daf-*2 over hermaphrodite WTs (P<0.0001 each). Male *daf-2* overall survival was significantly improved over WT males (logrank P<0.0001), and hazard ratio was significantly reduced (HR=0.0185; P<0.0001). Survival at the 75^th^ and 90^th^ percentiles of life were significantly improved in male *daf-*2 over male WTs (P<0.0001 each). Interestingly, male WT overall survival was significantly improved over hermaphrodite WT (logrank P<0.0001) and hazard ratio was significantly reduced (HR=0.4424; P<0.0001). Survival at the 75^th^ and 90^th^ percentiles of life were significantly improved in male WTs over hermaphrodite WTs (P<0.0001 each). Male *daf-2* overall survival was significantly improved over hermaphrodite *daf-2* survival (logrank P<0.0001) and hazard ratio was significantly reduced (HR=0.0585; P<0.0001). Survival at the 75^th^ and 90^th^ percentiles of life were significantly improved in male *daf-*2 over hermaphrodite *daf-2* (P<0.0001 each). Results for statistical analyses of the pooled survival analyses are presented in Table 1.

**Table 1.** Survival statistics for the aggregated lifespan data from Figure 1. Note these are visualized in Figure S1.

| Group | Median Lifespan | % Median Difference | Mean Lifespan | % Mean Difference | Logrank p-value | Cox Proportional Hazard |  | Maximal Lifespan |  |
| --- | --- | --- | --- | --- | --- | --- | --- | --- | --- |
|  |  |  |  |  |  | Hazard Ratio | p-value | 75th p-value | 90th p-value |
| ♀ WT (n=185) | 14 days | 103.6 | 14.7 days | 153.1 | <0.0001 | 1 | <0.0001 | <0.0001 | <0.0001 |
| ♀ daf-2(e1270) (n=132) | 14.5 days |  | 22.5 days |  |  | 0.4753 |  |  |  |
| ♂ WT (n=325) | 18 days | 408.3 | 19.4 days | 360.3 | <0.0001 | 1 | <0.0001 | <0.0001 | <0.0001 |
| ♂ daf-2(e1370) (n=353) | 73.5 days |  | 69.9 days |  |  | 0.0185 |  |  |  |
| ♀ WT (n=185) | 14 days | 128.6 | 14.7 days | 132.0 | <0.0001 | 1 | <0.0001 | <0.0001 | <0.0001 |
| ♂ WT (n=325) | 18 days |  | 19.4 days |  |  | 0.4424 |  |  |  |
| ♀ daf-2(e1270) (n=132) | 14.5 days | 506.9 | 22.5 days | 310.7 | <0.0001 | 1 | <0.0001 | <0.0001 | <0.0001 |
| ♂ daf-2(e1370) (n=353) | 73.5 days |  | 69.9 days |  |  | 0.0585 |  |  |  |

**Figure 1.**
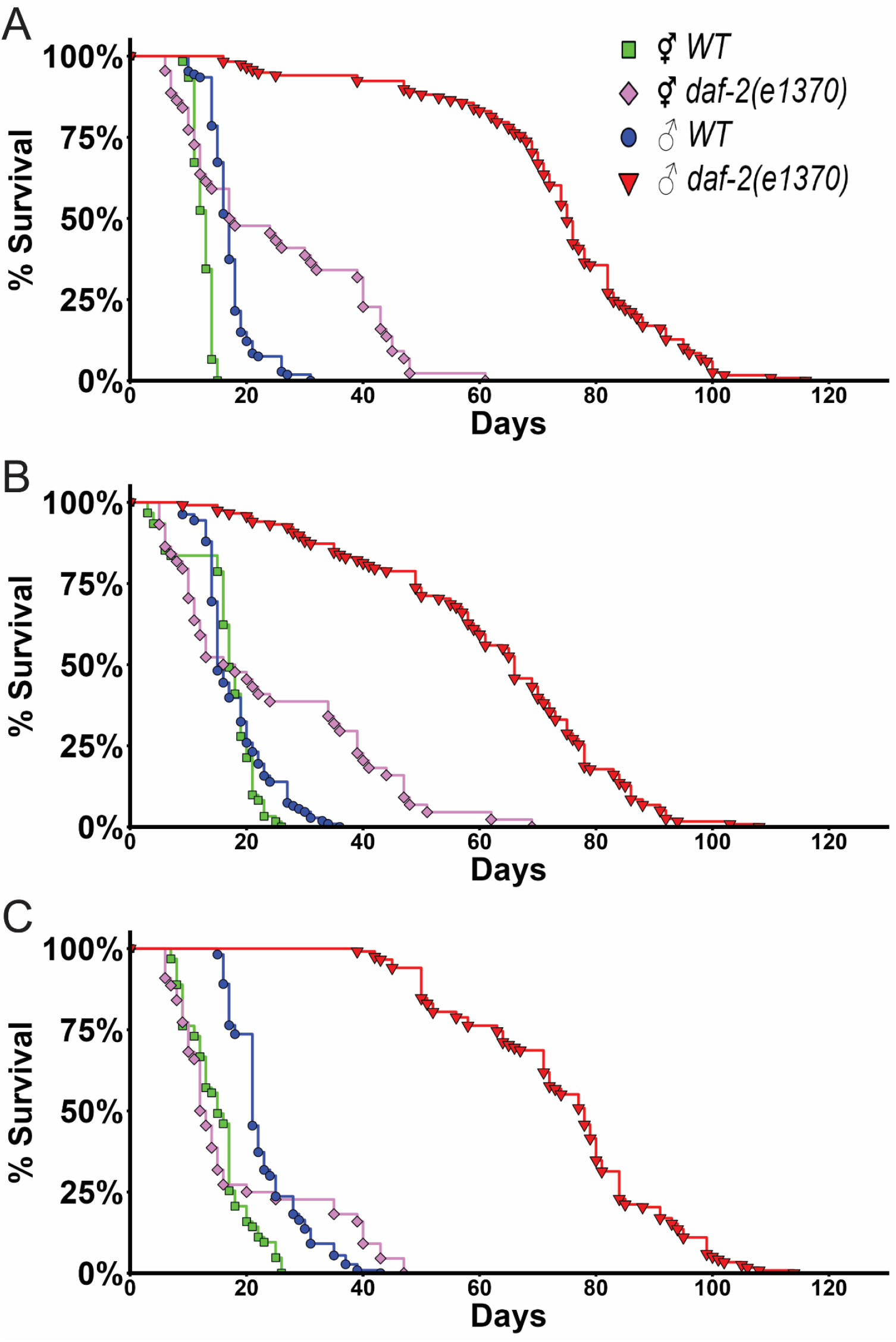
Loss of *daf-2* extends male lifespan fourfold. Kaplan-Meier survival curves for first (A), second (B) and third (C) independent replicates. N=44-118 per group in all panels. Statistical analysis is presented in Table 1.

### Male daf-2 Mutants are Longer and Accumulate More Lipids in Adulthood

Male *daf-2* mutants were observed to be more optically opaque than male WTs during adulthood. To investigate this observation, oil red o (ORO) staining was carried out to visualize neutral lipid accumulation in adult animals. Image analysis revealed a significant effect of genotype (P=0.0006) on body length, and a significant interaction effect of genotype X age (Px=0.0450) on body length. Post-hoc analyses of these data revealed male *daf-2* mutants were significantly longer than male WTs at day 14 of adulthood (Fig. 2A). Quantitative analysis of ORO staining revealed a significant effect of genotype (P<0.0001) and a significant interaction effect of genotype X age (Px<0.0001) on the percentage of the worm body stained with ORO. Post-hoc analyses of these data revealed that significantly less of the *daf-2* male body stained for ORO compared to WT males as day 1 adults, however at day 14 and day 20 *daf-2* males had significantly greater percentages of their body staining for ORO (Fig. 2B, 2D). Quantitative analysis of the mean ORO stain intensity revealed a significant effect of genotype (P<0.0001) and a significant interaction effect of genotype X age (Px<0.0001) on mean ORO intensity. Post-hoc analysis revealed significantly reduced ORO stain intensity compared to WT males at day 1, but significantly greater ORO stain intensity at day 14 and day 20 of adulthood (Fig. 2C, 2D). Together, these indicate that *daf-2* male *C. elegans* accumulate more lipids in adulthood.

**Figure 2.**
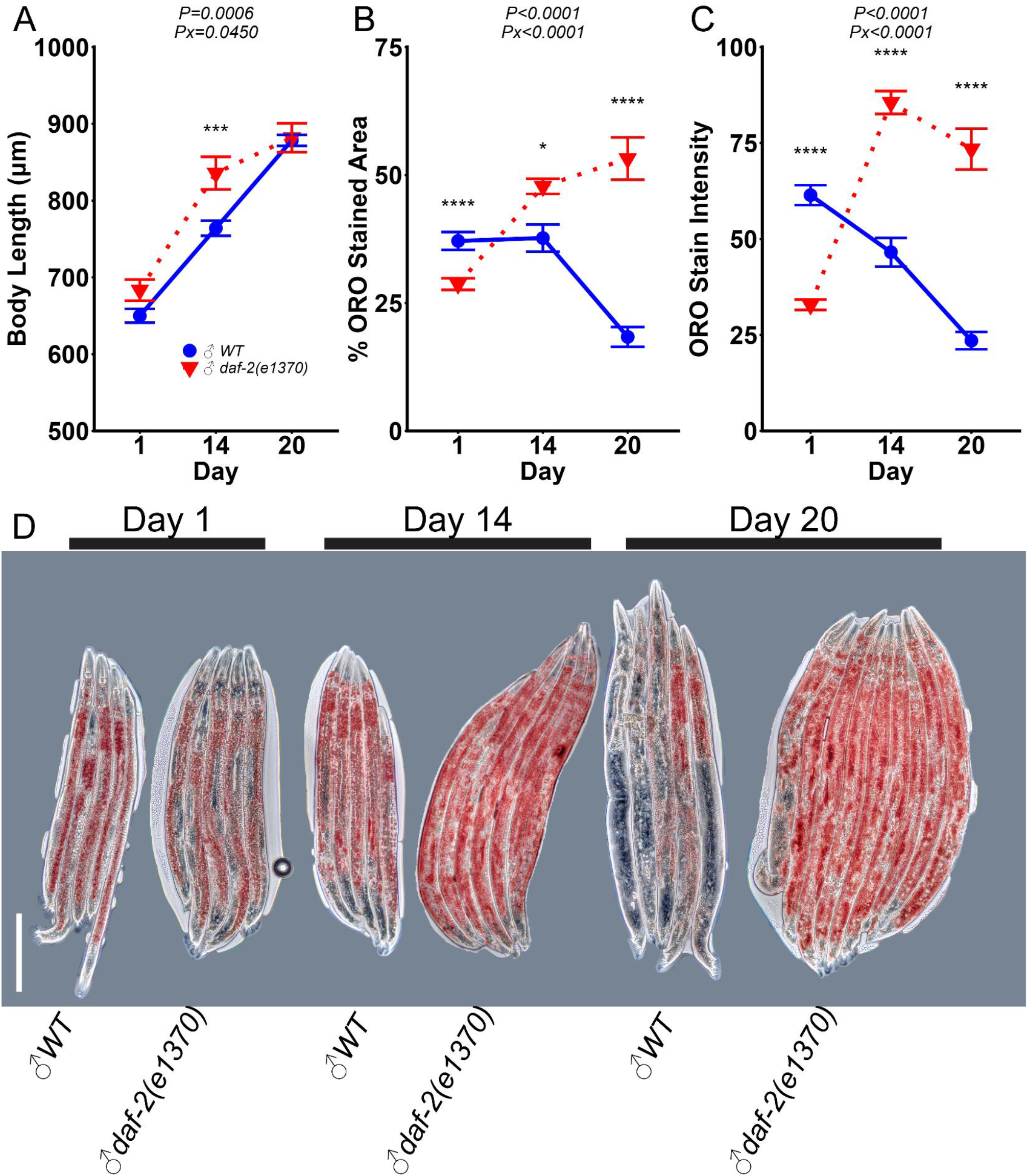
Long-lived *daf-2* mutant males accumulate and retain lipids in age. Quantification of body length (A). Quantification of the percentage of the worm body stained with oil red O (ORO) (B). Quantification of the average ORO stain intensity within the worm body (C). Representative images of stained worms (D). Scale bar represents 200µm. Statistical significance assessed by two-way ANOVA. Data presented as mean ± SEM. P represents the main effect of genotype; Px represents the genotype X age interaction effect. *P<0.05; ***P<0.001; ****P<0.0001 as determined by Tukey-HSD pairwise comparisons. N=13-41 per group.

### Male daf-2 Mutants Retain Motility and Resist Oxidative Stress

A key question regarding extreme lifespan extension is the preservation of health. Past work has demonstrated that the motility of *daf-2(e1370)* mutants is impaired, at least in in early hermaphrodite adulthood (Bansal et al., 2015; Gems et al., 1998; Mulcahy et al., 2013; Roy et al., 2022). This is undesirable, as it may represent periods of poor health. *daf-2(e1370)* hermaphrodites are known to resist oxidative stress, which is among the hypothesized mediators of their longevity (Bansal et al., 2015; Dues et al., 2019; Honda & Honda, 1999). We evaluated if the dramatically extended lifespan in our male *daf-2* mutants coincided with improvements in motility and stress resistance. Thrashing assay analysis revealed a significant effect of genotype (P<0.0001) and a significant interaction effect of genotype X age (Px<0.0001) on motility. Post-hoc analysis revealed that while thrashing rates were comparable at day 1 of adulthood, day 10 *daf-2* males displayed significantly greater thrashing rates compared to WTs (Fig. 3A). Hydrogen peroxide stress testing revealed that a significantly greater proportion of *daf-2* males survived 5mM and 10mM concentrations of hydrogen peroxide compared to WT males (P<0.0001 each; Fig. 3B). These data indicate that the male *daf-2* longevity is not simply a state of protracted frailty.

**Figure 3.**
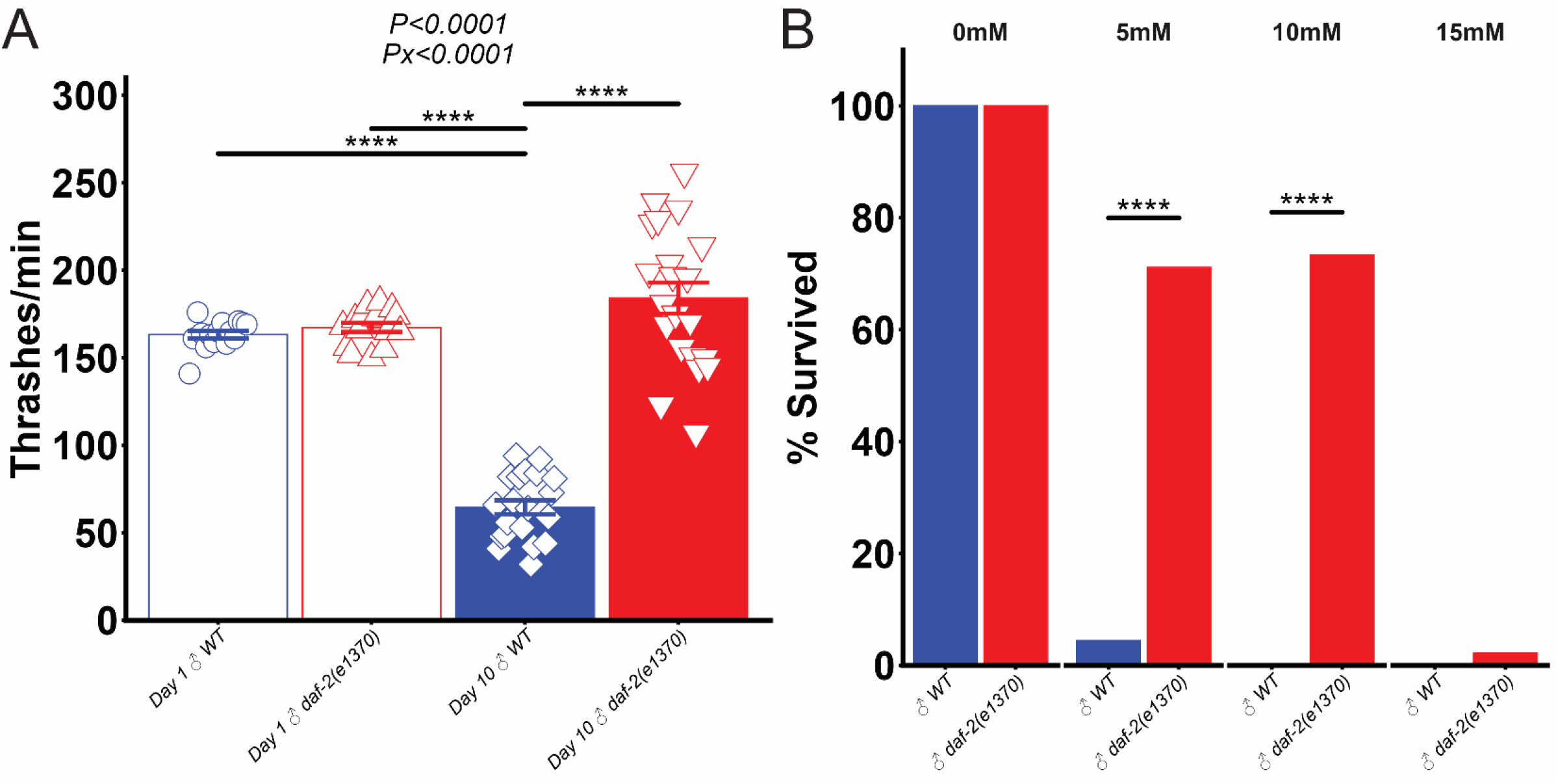
Long-lived *daf-2* mutant males retain motility and resist oxidative stress. Thrashing assay (A) and survival following a 4-hour incubation at the indicated hydrogen peroxide concentrations in day 1 males (B). Data presented as mean ± SEM with points representing individual animals (A) or as percentage surviving after incubation (B). P represents the main effect of genotype; Px represents the genotype X age interaction effect. ****P<0.0001 as determined by Tukey-HSD pairwise comparisons (A). ****P<0.0001 as determined by fisher exact test (B). N=15-21 per group (A) or N=27-46 per group (B).

### Functional DAF-16 is Required for the Extreme Male daf-2 Mutant Longevity

It has previously been demonstrated that improvements in wild-type male *C. elegans* longevity is not the result of reduced DAF-2 function (Gems & Riddle, 2000). It remains untested however, if the significant gains in male longevity conferred by *daf-2* mutation proceed through canonical signaling pathways or through multiple distinct mechanisms. To determine if the dramatic extension in male lifespan functions through canonical DAF-2-DAF-16 pathway as in hermaphrodites, we evaluated the lifespan of male *daf-2(e1370)/daf-16(mu86)* double mutants. The lifespan of male WTs, male *daf-16*, and male *daf-2/daf-16* mutants were generally comparable with each other and notably shorter than the male *daf-2* mutants (Fig. 4). Median lifespan of male *daf-16* mutants was 94.7% of the male WT median (18 days vs 19 days) and mean lifespan was 98.4% of the male WTs (18.9 days vs 19.2 days). Median lifespan of male *daf-2/daf-16* mutants was 84.2% of the male WTs (16 days vs 19 days) and mean lifespan was 93.2% of the male WTs (17.9 days vs 19.2 days). Median lifespan of male *daf-2/daf-16* mutants was 88.9% of the male *daf-16* mutant lifespan (16 days vs 18 days) and mean lifespan was 94.7% of the male *daf-16* mutants (17.9 days vs 18.9 days). Overall survival and hazard ratios were comparable in male *daf-16* mutants compared to male WTs (logrank P=0.3716, HR=1.1593, P=0.3310), but survival at the 90^th^ percentile of life was reduced (P=0.0029).

**Figure 4.**
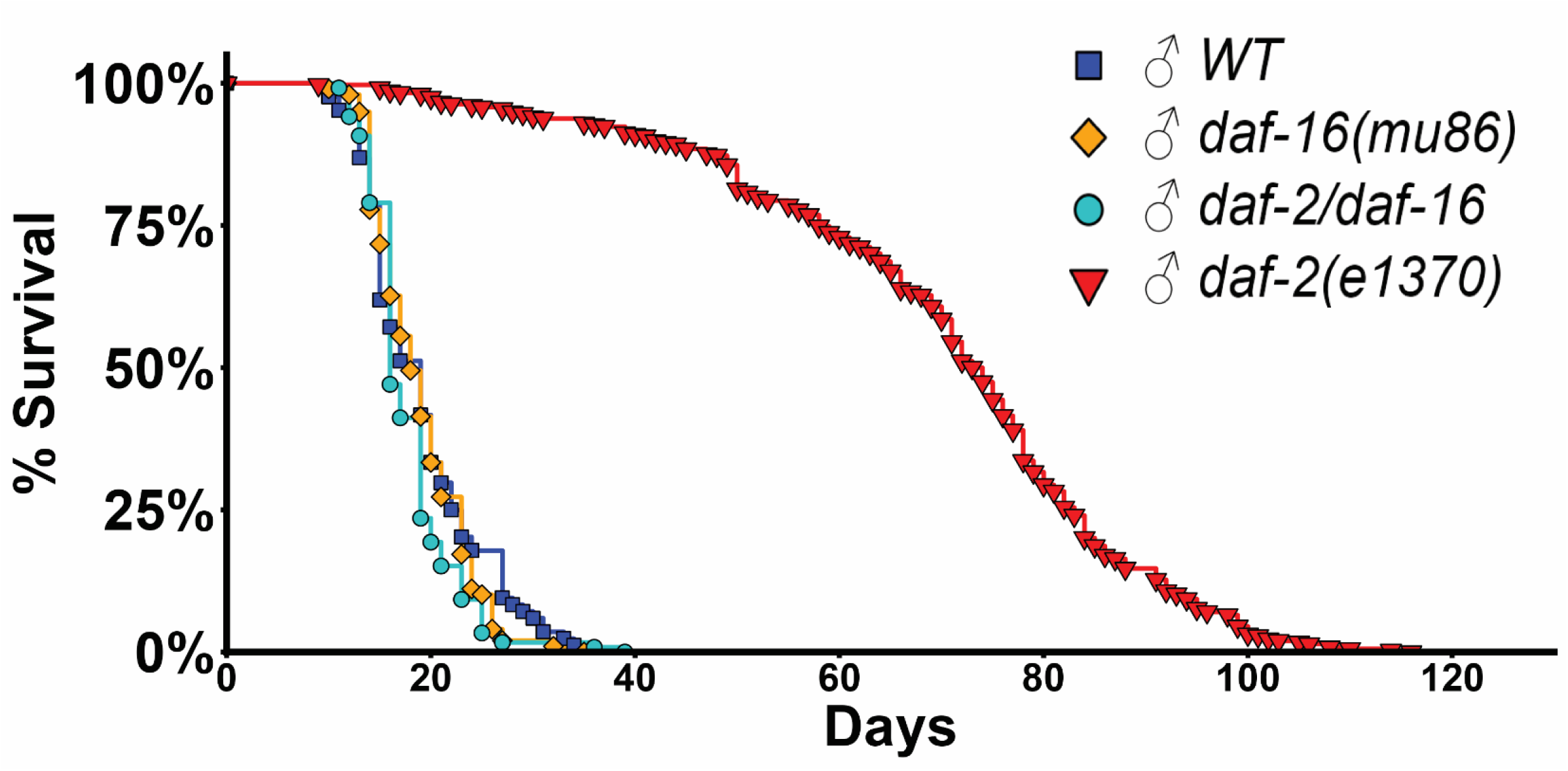
Functional *daf-16* is required for the extreme *daf-2* mutant male lifespan. Kaplan-Meier survival curves for WT, *daf-16(mu86)* and *daf-2(e1370) daf-16(mu86)* double mutant (*daf-2/daf-16*) males, as well as aggregated *daf-2(e1370)* male appended from figure 1 for visual reference. Statistical analysis is presented in Table 2.

Overall survival was not different in male *daf-16/daf-2* mutants compared to male WTs (logrank P=0.1877); however hazard ratio was significantly greater in male *daf-2/daf-16* mutants (HR=1.3443; P=0.0437). Survival at the 75^th^ percentile of life was reduced in male *daf-2/daf-16* mutants compared to male WTs (P=0.0322) and while a trend for reduced survival at the 90^th^ percentile was also observed, this failed to reach statistical significance (P=0.0883). Overall survival was not different between male *daf-2/daf-16* and male *daf-16* mutants, and while a trend for greater hazard ratios was observed in *daf-2/daf-16* mutants, this failed to reach statistical significance (P=0.0812). Survival at the 75^th^ percentile of life was reduced in male *daf-2/daf-16* mutants compared to male *daf-16* mutants (P=0.0203), but no differences were observed in the survival at the 90^th^ percentile of life. The results of statistical analyses are presented in Table 2. Together, these data indicate that the extreme lifespan of male *C. elegans* without functional DAF-2 requires functional DAF-16.

**Table 2.** Survival statistics for the lifespan data presented in Figure 4.

| Group | Median Lifespan | % Median Difference | Mean Lifespan | % Mean Difference | Logrank p-value | Cox Proportional Hazard |  | Maximal Lifespan |  |
| --- | --- | --- | --- | --- | --- | --- | --- | --- | --- |
|  |  |  |  |  |  | Hazard Ratio | p-value | 75th p-value | 90th p-value |
| ♂ WT (n=84) | 19.0 days | 94.7 | 19.2 days | 98.4 | 0.3716 | 1 | 0.3310 | 0.7395 | 0.0029 |
| ♂ daf-16(mu86) (n=99) | 18.0 days |  | 18.9 days |  |  | 1.1593 |  |  |  |
| ♂ WT (n=84) | 19.0 days | 84.2 | 19.2 days | 93.2 | 0.1877 | 1 | 0.0437 | 0.0322 | 0.0883 |
| ♂ daf-2/daf-16 (n=119) | 16.0 days |  | 17.9 days |  |  | 1.3443 |  |  |  |
| ♂ daf-16(mu86) (n=99) | 18.0 days | 88.9 | 18.9 days | 94.7 | 0.1877 | 1 | 0.0812 | 0.0203 | 0.6591 |
| ♂ daf-2/daf-16 (n=119) | 16.0 days |  | 17.9 days |  |  | 1.2713 |  |  |  |

## Discussion

Our current study reveals that the physiological plasticity inherent in the IIS signaling architecture is far more expansive than the benchmarks established over the past three decades. This work builds upon the seminal findings of Gems and Riddle, who first identified the sex-dependent lifespan effects of IIS and noted that *daf-2* mutant males possess a significant survival advantage over their wild-type counterparts (Gems & Riddle, 2000). These earlier reports focused primarily on the behavioral and environmental modulators of male lifespan such as the high energetic costs of mating and density-dependent stress. Our data here reveals a far more focused genetic expansion. By identifying a 110-day “longevity ceiling,” we demonstrate that the male IIS output is not merely a variation of the hermaphrodite program, but a distinct, fourfold amplification of somatic maintenance.

Interestingly, the 110-day longevity expansion is entirely dependent on functional DAF-16, yet it exhibits a striking “effect asymmetry.” In hermaphrodites, DAF-16 activation via loss of DAF-2 activity yields an approximately twofold lifespan extension (Kenyon et al., 1993; Ogg et al., 1997); in males, the same DAF-16 activation produces a nearly fourfold increase. This discrepancy suggests that DAF-16 operates within a unique male-specific chromatin environment, potentially binding to an entirely different network of “unknown targets.” This permissive chromatin landscape allows male DAF-16 to unlock latent survival pathways that remain dormant or suppressed in the hermaphrodite situation. In this vein, our work is consistent with previous reports that ablation of the *daf-2(e1370)* germ line precursor cells extends hermaphrodite *daf-2(e1370)* lifespan to a level comparable to our mutant males (Hsin & Kenyon, 1999). These benefits were dependent on functional DAF-16 and DAF-12, and thus it was proposed that the mechanism for this improvement was that the germline produces signals that impair DAF-16 and DAF-12 activity, shortening lifespan. Our data presented here build off this proposed mechanism and implicate oocytes as the origin for these signals.

Further, the physical foundation of this 4-fold longevity lies in a post-developmental “metabolic metamorphosis.” We observed a counter-intuitive lipid inversion: *daf-2* mutant males are not born with superior energy reserves—their lipid levels at Adult Day 1 are significantly lower than those of wild-type males. However, by Day 14 and Day 20, this phenotype completely reverses, with *daf-2* mutant males exhibiting massive neutral lipid accumulation. The loss of DAF-2 function has previously been associated with increased lipid accumulation (Kimura et al., 1997; Murphy et al., 2003), and impairments in *daf-2* mutant lipid accumulation results in a partial normalization of lifespan (Wang et al., 2008). The increased lipid accumulation through adulthood in our animals may indicate that a pro-longevity lipid profile contributes to the extended life of our animals. This suggests that IIS disruption in males triggers a critical temporal window of metabolic reprogramming, reallocating resources away from early-life energy expenditure and toward late-life somatic preservation.

Importantly, this 4-fold lifespan expansion challenges the canonical premise that extreme longevity requires a compensatory reduction in somatic growth. In classic models of aging, ranging from *daf-2* mutant hermaphrodites to dwarf mice and mammals subjected to caloric restriction, lifespan extension is usually linked to stunted growth or reduced physical size (Bartke, 2017). Here, we observe a surprising uncoupling of this growth-longevity axis that challenges the “disposable soma” theory. Rather than suffering the developmental penalties, as implied by the disposable soma theory (Kirkwood, 1977) and typical of these established longevity interventions, *daf-2* mutant males undergo a dynamic, age-progressive physical expansion. Quantitative imaging confirms a significant interaction between genotype and age driving this growth divergence, resulting in *daf-2* mutant males growing significantly longer than their wild-type counterparts by adult day 14. This temporal phenomenon, in which 110-day survival coexists with continuous late-life somatic enlargement, dictates a radical reallocation of resources unseen in traditional longevity interventions.

In summary, the male-specific DAF-16/FOXO network acts as a hidden “multiplier,” unlocking a survival potential previously unobserved in metazoans. This provides not only a new blueprint for sex-specific biomarkers of aging but also identifies the maximal plastic limits of healthy lifespan extension through the allosteric modulation of core signaling pathways.

## Methods

### Strains

The original N2 (WT) animals were a gift from the laboratory of Dr. Haoseng Sun at the University of Alabama at Birmingham. *daf-2(e1370)* (strain #: CB1370) and *daf-16(mu86)* (strain #: CF1038) mutants were acquired from the Caenorhabditis Genetics Center at the University of Minnesota. Double mutant *daf-2(e1370)*;*daf-16(mu86)* animals were generated by mating *daf-2(e1370)* males (generated as detailed below) with daf-16 hermaphrodites for 48 hours.

Successful transmission of the *daf-2(e1370)* mutation was confirmed by PCR with forward primer (5’-3’) aatccgtaaggcagatgacg and the reverse primer (5’-3’) cgtaaggacttgtacgccaa, followed by a BlpI (New England Biolabs) digestion carried out according to the manufacturer suggested protocol. Successful transmission of the *daf-16(mu86)* mutation was confirmed by PCR with the common forward primer (5’-3’) tcgccttcatcatctatcccc, the WT reverse primer (5’-3’) gatgggggcaatctgaggt, and the mutant reverse primer (5’-3’) tcactatctcttacctttgtagtcgt.

### Worm maintenance and Survival Analysis

Male *daf-2(e1370)* mutants were generated by picking spontaneous males from our *daf-2(e1370)* colony and mating them to hermaphrodites to increase the frequency of males in subsequent progeny. Male *daf-16(mu86)* and *daf-2(e1370);daf-16(mu86)* mutants were generated by incubating L4 animals at 30°C for 5-6 hours to increase the frequency of males in subsequent progeny. Animals were grown at 15°C until L4 stage, then shifted to 20°C for survival studies. Animals were maintained on standard NGM plates with palmitic acid barriers for all assays to prevent animals from leaving the plate (Beydoun et al., 2024) and desiccating on the plate walls. Hermaphrodites were transferred to new plates every other day until egg-laying ceased and then every three days after that (Park et al., 2017). The males were transferred every three days as well.

### Oil Red O staining

We employed a previously described protocol(Stuhr et al., 2022) with minor modifications. A stock solution of Oil Red O was prepared by dissolving 0.25g Oil Red O into 49.75ml isopropanol followed by vacuum filtration. A working solution was prepared before staining by diluting the stock solution to a concentration of 60% (v/v) in water. This working solution was agitated overnight in the dark at room temperature to dissolve any aggregates. Animals at the indicated ages were washed off their plates with 1x phosphate buffered saline with 0.01% (v/v) Triton X-100 (PBST), transferred to 1.5ml microcentrifuge tubes, and pelleted by centrifugation. The supernatant was removed and the washing/pelleting steps were repeated two additional times. Washed pellets were incubated in 0.9ml isopropanol at room temperature with agitation for three minutes, pelleted by centrifugation, and then the supernatant was removed. 1ml prepared working Oil Red O staining solution was added to the pellet, and then then agitated in the dark at room temperature for 2 hours. Animals were pelleted, the supernatant was removed and incubated in PBST for 30 minutes at room temperature with gentle agitation. This was repeated for 4 total washes. Stained animals were positioned on a 5% agar pad, oriented with an eyelash pick, and mounted on microscope slides. Brightfield tile scans of whole worms were acquired using a Leica DMI8 microscope with a Leica DMC2900 camera and motorized stage. Images were color deconvoluted in Fiji(Schindelin et al., 2012) using color deconvolution 1.7 with FastRed FastBlue DAB vectors. The resulting red channels were inverted and subjected to Otsu thresholding without manual adjustment. Regions of interest were manually drawn around the worm bodies and area fraction and mean grey values were used to compare percentage of the worm body stained with Oil Red O and the intensity of the Oil Red O staining, respectively.

### Thrashing assay

Approximately five worms were placed in 20uL of M9 buffer and allowed to acclimate for a few seconds to the solution before recording video acquisition of body bending movements for one minute. For counting body-bending, the video was slowed down to half speed (0.5x) and movements were recorded by an experimenter blinded to the animal’s group membership. Recordings were collected using a Nikon AZ100M dissecting scope with a Nikon DS-Fi2 camera.

### Hydrogen Peroxide Stress Assay

We followed a previously published protocol(Huang et al., 2022) with minor modifications. Between 9 and 16 synchronized day 1 adults were transferred into each well of a 12-well plate containing S-basal buffer supplemented with various concentrations of H_2_O_2_ (0mM, 5mM, 10mM, and 15mM). Plates were incubated at 20°C for 4 hours. Following incubation 100μl of 1 mg/ml catalase was transferred to neutralize the H_2_O_2_. The mortality of worms was recorded following neutralization.

### Statistical analysis

Lifespan data were visualized using Kaplan-Meier survival curves. Overall survival was statistically compared using logrank tests, and hazard ratios were compared using cox proportional hazard testing. Maximal lifespan was evaluated by comparing the proportion of individuals within each group remaining alive at the 75^th^ and 90^th^ percentiles of survival using the quantile-regression approach previously described(Wang et al., 2004). Where group means were compared, a two-way ANOVA was used with Tukey HSD post-hoc tests carried out where significant main effects or interaction effects were detected. Where proportions were compared, a fisher exact test was used. For all statistical tests significance was established at P<0.05. All analyses were carried out and all figures were generated using the R programming language (R Core Team, 2025).

## Contributions

Liou Y. Sun and Steven N. Austad conceptualized the study. Liou Y. Sun oversaw overall direction and secured funding. Mike Russell, Michelle Lin, and Liou Y. Sun designed experiments. Mike Russell and Michelle Lin conducted experiments and collected data. Alexander Tate Lasher analyzed data. Mike Russell took the lead in writing the manuscript. Mike Russell, Alexander Tate Lasher, Michelle Lin, Liou Y. Sun, and Steven N. Austad edited the manuscript. All authors provided critical feedback that helped shape the research, analysis, and final report presented here.

## Acknowledgements

The authors would also like to recognize the critical feedback and insightful comments from all members of the Sun Lab that helped conceive this manuscript.

## Data availability

All data used to generate the statistical analyses and figures in this manuscript are available from the corresponding author upon reasonable request.

## References

Austad, S. N., & Fischer, K. E. (2016). Sex Differences in Lifespan. Cell Metab, 23(6), 1022–1033. 10.1016/j.cmet.2016.05.019

Bansal, A., Zhu, L. J., Yen, K., & Tissenbaum, H. A. (2015). Uncoupling lifespan and healthspan in Caenorhabditis elegans longevity mutants. Proc Natl Acad Sci U S A, 112(3), E277–286. 10.1073/pnas.1412192112

Bartke, A. (2017). Somatic growth, aging, and longevity. NPJ Aging Mech Dis, 3, 14. 10.1038/s41514-017-0014-y

Beydoun, S., Sridhar, A., Bhandari, M., Kitto, E. S., & Leiser, S. F. (2024). Polyethylene glycol as an improved barrier to prevent fleeing in C. elegans. MicroPubl Biol, 2024. 10.17912/micropub.biology.001288

Bokov, A. F., Garg, N., Ikeno, Y., Thakur, S., Musi, N., DeFronzo, R. A., Zhang, N., Erickson, R. C., Gelfond, J., Hubbard, G. B., Adamo, M. L., & Richardson, A. (2011). Does reduced IGF-1R signaling in Igf1r+/mice alter aging? PLoS One, 6(11), e26891. 10.1371/journal.pone.0026891

Clancy, D. J., Gems, D., Harshman, L. G., Oldham, S., Stocker, H., Hafen, E., Leevers, S. J., & Partridge, L. (2001). Extension of life-span by loss of CHICO, a Drosophila insulin receptor substrate protein. Science, 292(5514), 104–106. 10.1126/science.1057991

Dues, D. J., Andrews, E. K., Senchuk, M. M., & Van Raamsdonk, J. M. (2019). Resistance to Stress Can Be Experimentally Dissociated From Longevity. J Gerontol A Biol Sci Med Sci, 74(8), 1206–1214. 10.1093/gerona/gly213

Gems, D., & Riddle, D. L. (2000). Genetic, behavioral and environmental determinants of male longevity in Caenorhabditis elegans. Genetics, 154(4), 1597–1610. 10.1093/genetics/154.4.1597

Gems, D., Sutton, A. J., Sundermeyer, M. L., Albert, P. S., King, K. V., Edgley, M. L., Larsen, P. L., & Riddle, D. L. (1998). Two pleiotropic classes of daf-2 mutation affect larval arrest, adult behavior, reproduction and longevity in Caenorhabditis elegans. Genetics, 150(1), 129–155. 10.1093/genetics/150.1.129

Holzenberger, M., Dupont, J., Ducos, B., Leneuve, P., Géloën, A., Even, P. C., Cervera, P., & Le Bouc, Y. (2003). IGF-1 receptor regulates lifespan and resistance to oxidative stress in mice. Nature, 421(6919), 182–187. 10.1038/nature01298

Honda, Y., & Honda, S. (1999). The daf-2 gene network for longevity regulates oxidative stress resistance and Mn-superoxide dismutase gene expression in Caenorhabditis elegans. FASEB J, 13(11), 1385–1393.

Hsin, H., & Kenyon, C. (1999). Signals from the reproductive system regulate the lifespan of C. elegans. Nature, 399(6734), 362–366. 10.1038/20694

Huang, M., Hong, M., Hou, X., Zhu, C., Chen, D., Chen, X., Guang, S., & Feng, X. (2022). H3K9me1/2 methylation limits the lifespan of daf-2 mutants in C. elegans. Elife, 11. 10.7554/eLife.74812

Kenyon, C. (2011). The first long-lived mutants: discovery of the insulin/IGF-1 pathway for ageing. Philos Trans R Soc Lond B Biol Sci, 366(1561), 9–16. 10.1098/rstb.2010.0276

Kenyon, C., Chang, J., Gensch, E., Rudner, A., & Tabtiang, R. (1993). A C. elegans mutant that lives twice as long as wild type. Nature, 366(6454), 461–464. 10.1038/366461a0

Kimura, K. D., Tissenbaum, H. A., Liu, Y., & Ruvkun, G. (1997). daf-2, an insulin receptor-like gene that regulates longevity and diapause in Caenorhabditis elegans. Science, 277(5328), 942–946. 10.1126/science.277.5328.942

Kirkwood, T. B. (1977). Evolution of ageing. Nature, 270(5635), 301–304. 10.1038/270301a0

MacNee, W., Rabinovich, R. A., & Choudhury, G. (2014). Ageing and the border between health and disease. Eur Respir J, 44(5), 1332–1352. 10.1183/09031936.00134014

Mulcahy, B., Holden-Dye, L., & O’Connor, V. (2013). Pharmacological assays reveal agerelated changes in synaptic transmission at the Caenorhabditis elegans neuromuscular junction that are modified by reduced insulin signalling. J Exp Biol, 216(Pt 3), 492–501. 10.1242/jeb.068734

Murphy, C. T., McCarroll, S. A., Bargmann, C. I., Fraser, A., Kamath, R. S., Ahringer, J., Li, H., & Kenyon, C. (2003). Genes that act downstream of DAF-16 to influence the lifespan of Caenorhabditis elegans. Nature, 424(6946), 277–283. 10.1038/nature01789

Ogg, S., Paradis, S., Gottlieb, S., Patterson, G. I., Lee, L., Tissenbaum, H. A., & Ruvkun, G. (1997). The Fork head transcription factor DAF-16 transduces insulin-like metabolic and longevity signals in C. elegans. Nature, 389(6654), 994–999. 10.1038/40194

Park, H. H., Jung, Y., & Lee, S. V. (2017). Survival assays using Caenorhabditis elegans. Mol Cells, 40(2), 90–99. 10.14348/molcells.2017.0017

R Core Team. (2025). R: A Language and Environment for Statistical Computing. In R Foundation for Statistical Computing. https://www.R-project.org/

Roy, C., Molin, L., Alcolei, A., Solyga, M., Bonneau, B., Vachon, C., Bessereau, J. L., & Solari, F. (2022). DAF-2/insulin IGF-1 receptor regulates motility during aging by integrating opposite signaling from muscle and neuronal tissues. Aging Cell, 21(8), e13660. 10.1111/acel.13660

Schindelin, J., Arganda-Carreras, I., Frise, E., Kaynig, V., Longair, M., Pietzsch, T., Preibisch, S., Rueden, C., Saalfeld, S., Schmid, B., Tinevez, J. Y., White, D. J., Hartenstein, V., Eliceiri, K., Tomancak, P., & Cardona, A. (2012). Fiji: an open-source platform for biological-image analysis. Nat Methods, 9(7), 676–682. 10.1038/nmeth.2019

Stuhr, N. L., Nhan, J. D., Hammerquist, A. M., Van Camp, B., Reoyo, D., & Curran, S. P. (2022). Rapid Lipid Quantification in Caenorhabditis elegans by Oil Red O and Nile Red Staining. Bio Protoc, 12(5), e4340. 10.21769/BioProtoc.4340

Suh, Y., Atzmon, G., Cho, M. O., Hwang, D., Liu, B., Leahy, D. J., Barzilai, N., & Cohen, P. (2008). Functionally significant insulin-like growth factor I receptor mutations in centenarians. Proc Natl Acad Sci U S A, 105(9), 3438–3442. 10.1073/pnas.0705467105

Tatar, M., Kopelman, A., Epstein, D., Tu, M. P., Yin, C. M., & Garofalo, R. S. (2001). A mutant Drosophila insulin receptor homolog that extends life-span and impairs neuroendocrine function. Science, 292(5514), 107–110. 10.1126/science.1057987

Wang, C., Li, Q., Redden, D. T., Weindruch, R., & Allison, D. B. (2004). Statistical methods for testing effects on “maximum lifespan”. Mech Ageing Dev, 125(9), 629–632. 10.1016/j.mad.2004.07.003

Wang, M. C., O’Rourke, E. J., & Ruvkun, G. (2008). Fat metabolism links germline stem cells and longevity in C. elegans. Science, 322(5903), 957–960. 10.1126/science.1162011

